# Comparison of target enrichment strategies for ancient pathogen DNA

**DOI:** 10.1101/2020.07.09.195065

**Authors:** Anja Furtwängler, Judith Neukamm, Lisa Böhme, Ella Reiter, Melanie Vollstedt, Natasha Arora, Pushpendra Singh, Stewart T. Cole, Sascha Knauf, Sébastien Calvignac-Spencer, Ben Krause-Kyora, Johannes Krause, Verena J. Schuenemann, Alexander Herbig

**Author notes:** These authors jointly supervised this study (Verena J. Schuenemann, Alexander Herbig).

## Abstract

In ancient DNA research, the degraded nature of the samples generally results in poor yields of highly fragmented DNA, and targeted DNA enrichment is thus required to maximize research outcomes. The three commonly used methods – (1) array-based hybridization capture and in-solution capture using either (2) RNA or (3) DNA baits – have different characteristics that may influence the capture efficiency, specificity, and reproducibility. Here, we compared their performance in enriching pathogen DNA of *Mycobacterium leprae* and *Treponema pallidum* of 11 ancient and 19 modern samples. We find that in-solution approaches are the most effective method in ancient and modern samples of both pathogens, and RNA baits usually perform better than DNA baits.

**Method summary:** We compared three targeted DNA enrichment strategies used in ancient DNA research for the specific enrichment of pathogen DNA regarding their efficiency, specificity, and reproducibility for ancient and modern *Mycobacterium leprae* and *Treponema pallidum* samples. Array-based capture and in-solution capture with RNA and DNA baits were all tested in three independent replicates.

## Main Text

The field of ancient DNA (aDNA), which studies DNA retrieved from paleontological and archaeological material, was revolutionized by the invention of high-throughput sequencing (HTS). In combination with HTS, the development of targeted DNA enrichment protocols has made a crucial contribution in advancing aDNA research during the last decade.

As DNA decays over time, aDNA is usually only present in trace amounts of highly fragmented sequences **(1, 2, 3)**. Detecting endogenous pathogen aDNA from archaeological material is additionally compounded by the larger amount of background DNA from the environment including soil microorganisms. Furthermore, the background of host DNA in ancient remains is an additional obstacle in order to obtain ancient pathogen DNA. Shotgun sequencing of libraries from aDNA extracts to sufficient genomic coverage is, therefore, cost-intensive **(4)**. To circumvent this problem, specific regions of interest such as bacterial chromosomes, mammalian mitochondrial genomes, or regions with single-nucleotide-polymorphisms (SNP) are often target-enriched before sequencing **(4)**. Aside from its application in aDNA sequencing, targeted DNA enrichment is also useful to retrieve pathogen DNA from clinical samples, particularly for infectious agents that are found in low quantities in the host organism and which are difficult to culture, as is the case for *Mycobacterium leprae* and *Treponema pallidum.* Removal of background DNA prior to sequencing increases the yield of pathogen DNA, and thus allows valuable information for epidemiologists investigating outbreaks to be obtained.

For the enrichment of entire bacterial and mammalian chromosomes, there are currently three methods available, which are based on hybridization capture **(5)**: DNA microarrays (here represented by SureSelect from Agilent Technologies), in-solution capture with DNA baits (represented by SureSelect from Agilent Technologies according to Fu and colleagues **(6)**) and in-solution capture with RNA baits (here represented by myBaits® from Arbor Biosciences).

In the case of the DNA array-based method, up to a million artificial DNA baits are printed on the surface of a glass slide **(7)**. Additionally, there is the possibility to perform in-solution capture with baits cleaved from the glass slides and used right away or immortalized in DNA bait libraries **(6)**. The second in-solution approach uses up to 100,000 artificial RNA baits. The three approaches rely on the hybridization of target fragments to the complementary sequence of the baits (immobilized or in-solution), which can be levered to wash background DNA away.

To date there has been to our knowledge, no statistical comparison of the performance of all three methods: microarrays, in-solution capture with DNA baits, and in-solution capture with RNA baits **(6)**. So far only microarrays and the in-solution capture with DNA baits were compared for *Salmonella enterica* and no replicates for statistical assessment were produced **(8)**.

Here, we present results from the enrichment of modern and ancient samples containing pathogen DNA, using the three aforementioned approaches. All samples had previously tested positive but had also shown low amounts of target DNA for *M. leprae* or *T. pallidum* (Supplementary Table 1).

The different enrichment concepts tested were chosen to represent methods as they are applied in ongoing research and therefore not only differ in the technology used (DNA *vs.* RNA baits, immobilized *vs.* in-solution) but also in the design such as bait length and number of unique baits, which might have an effect on the performance.

We used eight ancient samples positive for *M. leprae* and six modern libraries from leprosy patients that were shown to contain *M. leprae* DNA (Supplementary Note 1). Genetic data from the ancient and modern *M. leprae* samples were previously published in **9** and **10**. Samples with less than 0.6 % endogenous bacterial DNA were selected.

Modern *T. pallidum* samples (n=13) were previously published in **12** and **13**. Three ancient extracts of *T. pallidum* were used from **14**. The portion of endogenous DNA for the selected *T. pallidum* samples was below 0,01 % for ancient and modern samples.

Starting from existing sequencing libraries all three methods were applied with three independent replicates each (see Figure. 1 and Supplementary Note 1 for a detailed description of the methods, the newly generated data is available at the Sequence Read Archive under the BioProject PRJNA645054). Following the manufacturer’s suggestion for libraries with low yields of target DNA, we performed two successive rounds of hybridization for all methods. To investigate the effectiveness of this procedure, we compared results from the first and second rounds for the in-solution capture with RNA baits. We then evaluated differences in efficiency, reproducibility, and specificity across the three approaches by calculating mean coverage, standard deviation of the mean coverage, enrichment factor (calculated by dividing the % of target DNA after enrichment by the % of target DNA in the shotgun data), and the % of the genome covered 5-fold or more after normalizing the data of each bacterial species to the same number of raw reads (Supplementary Tables 2, 3 & 5 and Supplementary Figures 1 & 2).

**Figure 1.**
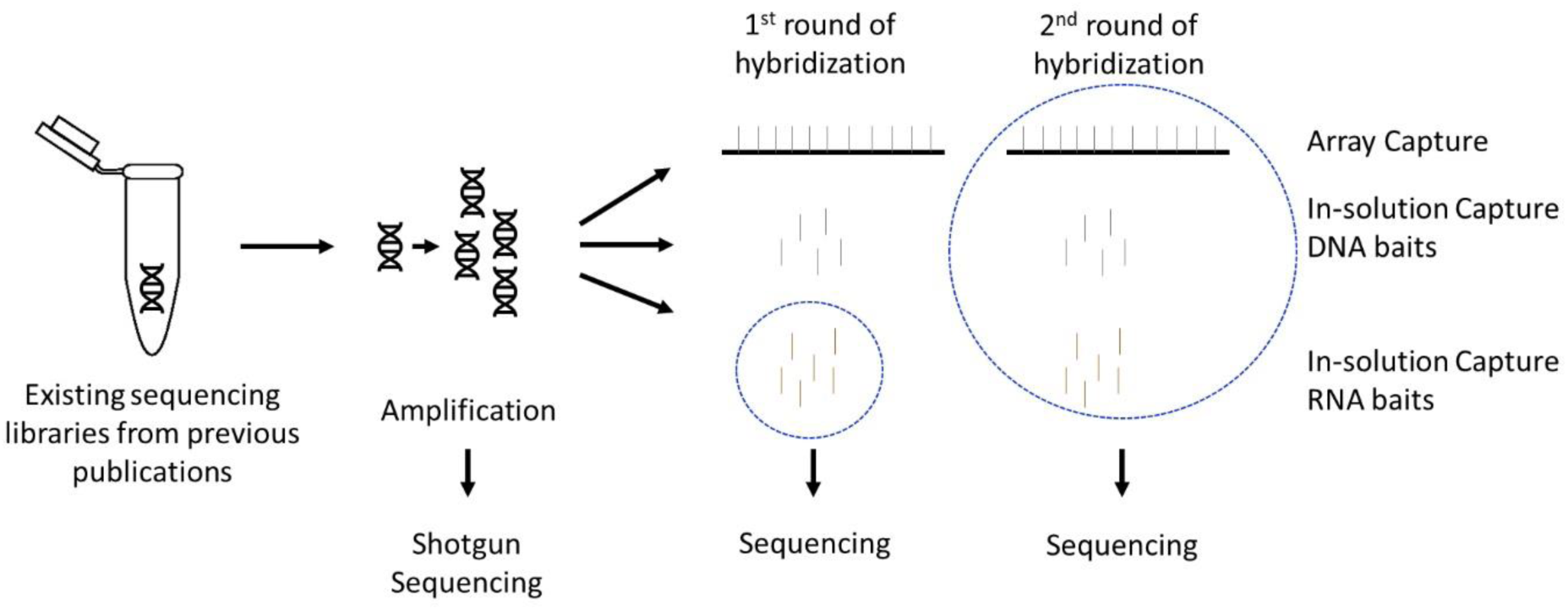
Schematic representation of the workflow. For all samples, the three different enrichment protocols were tested in three independent replicates. Blue circles indicate the libraries that were sequenced at each particular step.

For most ancient samples, the highest mean coverage (Figure 2A) is reached with the RNA bait in-solution capture (eight out of eleven, more details can be found in Supplementary Note 2 & 3, and SSupplementary Tables 1 & 2). On average the RNA bait capture results in a 1.5 and 20.0 times higher mean coverage than the DNA bait or the array capture, respectively. As illustrated in Figure. 2B, the highest enrichment factor is obtained in the RNA bait capture of ancient *T. pallidum* DNA (all three samples) and *M. leprae* (four samples showed best results for the RNA bait, three for the DNA bait, and one for the array), with values between 2-150x higher, compared to the other two approaches. An in-solution approach seems, therefore, to be advantageous for enriching ancient pathogen DNA.

**Figure 2.**
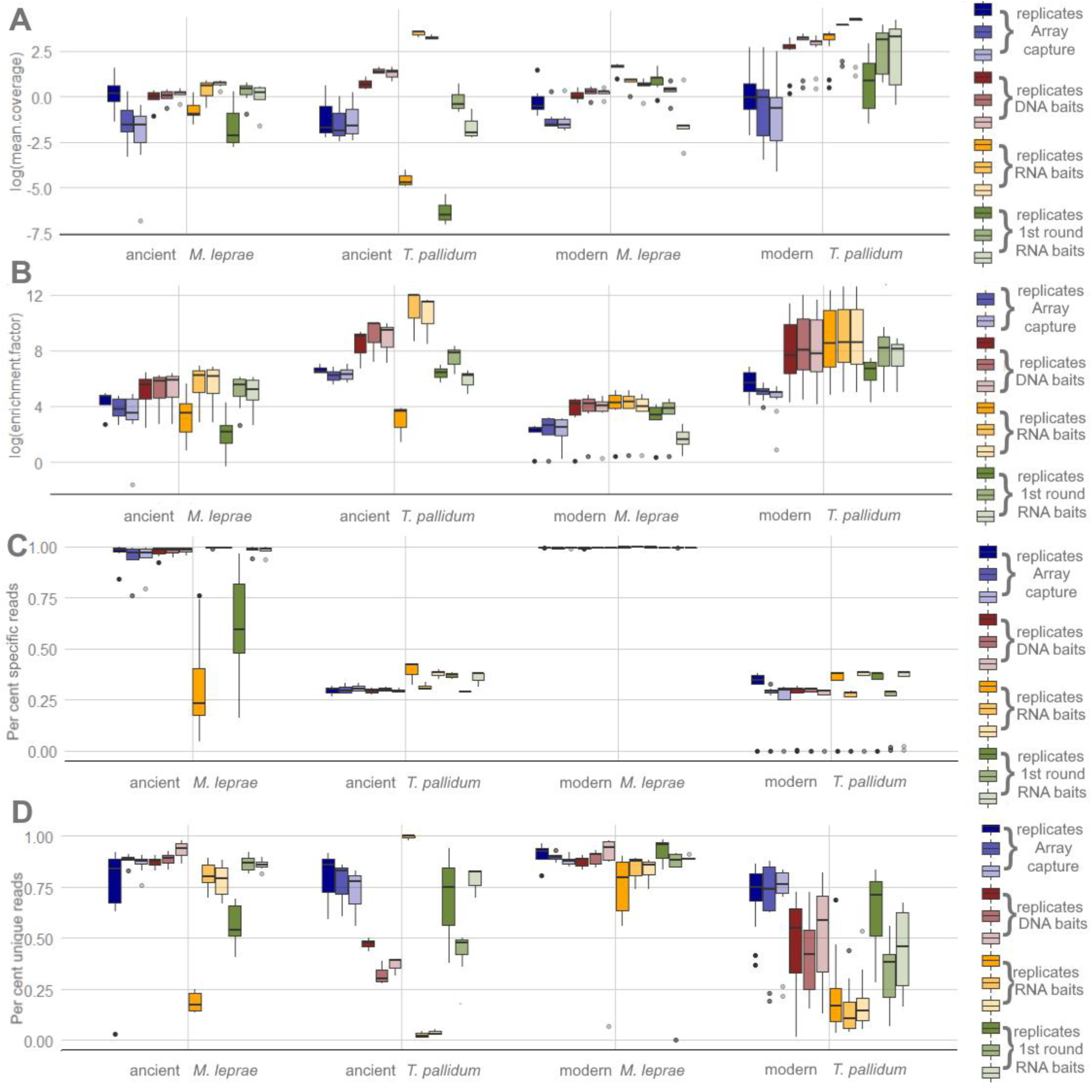
Differences between the three tested protocols in ancient and modern *M. leprae* and *T. pallidum* samples. A) Log-transformed values of the mean coverage. B) log-transformed values of the enrichment factor calculated by dividing the percentage of endogenous DNA by the percentage of endogenous DNA after shotgun sequencing. C) The proportion of specific reads corresponding to *M. leprae* and *T. pallidum* compared to other mycobacterial and treponemal reads, respectively. D) Percentage of unique reads calculated by the number of unique reads divided by the total number of sequences mapped to represent library complexity in *M. leprae* and *T. pallidum* samples.

A similar pattern can be observed in the data of the modern *M. leprae* and *T. pallidum* samples (Figures. 2A and 2B) further highlighting the performance of the in-solution approach in general and RNA baits in particular.

In-solution capture with DNA baits was used with robot-assistance in this study whereas the in-solution capture with RNA baits was performed in two different labs. Unsurprisingly, the DNA bait capture showed the smallest differences (2-to 50-fold lower) between the replicates whereas the RNA bait capture showed the largest and the DNA array capture was intermediate. Consistent conditions are therefore crucial for reproducibility.

Another important feature of targeted enrichment is specificity. We estimated the specificity of the three tested methods by comparing the number of reads specific to either *M. leprae* or *T. pallidum* in comparison to general mycobacterial or treponemal reads, respectively (Figure 2 C). Here, differences between the two pathogens can be observed. In the ancient and modern *T. pallidum* samples, the RNA bait capture consistently shows the highest proportion (up to 1.5 times higher) of specific reads. The same trend was observed for the libraries prepared from recent leprosy patient samples, i.e. modern samples of *M. leprae*. Only for ancient *M. leprae* samples, the DNA bait capture is more specific. The highest percentages of specific reads are not necessarily found in samples with high percentages of endogenous DNA in the shotgun data before enrichment.

For ancient and modern samples, due to high efficiency, reproducibility and specificity in-solution approaches are highly recommendable.

Two rounds of hybridization are routinely performed in aDNA research, which is expected to improve enrichment but may also reduce data complexity in terms of portions of unique reads. To formally investigate the effect of the second round of capture, we also sequenced the libraries only enriched with one round of hybridization with the RNA baits and compared the results to the second round of hybridization. The second round of hybridization resulted in an increase in the enrichment factor for ancient and modern *M. leprae* samples (with an average of 2x increase) as well as for *T. pallidum* samples (with an average of 17x increase), demonstrating the utility of such a second round of hybridization capture (Supplementary Table 5). On the other hand, when comparing the library complexity (Figure. 2 D and Supplementary Note 2 & 3, Supplementary Figure 3), we found a substantial loss of complexity after the second round of hybridization in all modern and ancient samples. This loss was reflected in the higher percentage of unique reads in all the reads mapped after the first round. Therefore, if the portion of endogenous DNA in a sample is high in the beginning it may be worthwhile considering whether a single round of capture combined with deeper sequencing is sufficient or even advantageous.

The three protocols also differ in terms of cost and effort. The most cost-intensive is the array-capture approach (∼673 € per sample), which requires additional equipment that is not usually necessary with the other approaches. The in-solution capture with DNA baits is, by contrast, cheaper once the baits are cleaved from the glass slide (∼56,23€ per sample), but the version that can be used for the immortalization of the baits by transforming them into a library is not freely available. The in-solution capture with RNA baits is more comparable to the DNA bait capture than to the array with ∼109 € per sample and it also needs the lowest number of additional equipment and reagents (Supplementary Table 7).

After a detailed comparison of the three tested methods it can be concluded that for ancient and modern pathogen samples, the RNA bait capture with two rounds of hybridization seems to be the most suitable. The generally high performance of the in-solution approach (mainly the one with RNA baits) for both bacterial species suggests that the findings are highly representative and comparable performance is also expected for a variety of other bacterial/microbial organisms.

## Author contributions

V.J.S., A.H. and J.K. conceived of the study. B.K. and S.C-S. provided RNA baits and sequencing libraries. N.A., P.S., S.T.C., S.K. provided sequencing libraries. A.F., L.B., E.R., M.V. performed the laboratory work. A.F. and J.N. performed the data analysis. A.F. and A.H. conducted the statistical analysis. A.F. designed the figures. A.F., V.J.S, and A.H. wrote the manuscript with input from all authors. All authors reviewed the manuscript.

## Acknowledgments

We thank all our colleagues providing samples for our study: Sarah Inskip (University of Cambridge, UK), Helen Donoghue (University College London, UK), Rodrigo Barquera (Max Planck Institute for the Science of Human History, Germany), Michael Taylor (University of Surrey, UK), Thomas Mendum (University of Surrey, UK), Graham Stewart (University of Surrey, UK), Simon Roffey (The Magdalen Hill Archaeological Research Project (MHARP) Winchester, UK), Phil Marter (The Magdalen Hill Archaeological Research Project (MHARP) Winchester, UK), Katie Tucker (Deutsches Archäologisches Institut, Berlin, Germany), Fabian Leendertz (Robert Koch Insitute, Berlin, Germany), Roman Wittig (Max Planck Institute for Evolutionary Anthropology, Leipzig, Germany), Anna Kjellström (University of Stockholm; Sweden), Christos Economou (University of Stockholm, Sweden), Petr Velemínský (National Museum, Czech Republic) Antónia Marcsik (University of Szeged, Hungary), Erika Molnár (University of Szeged, Hungary), György Pálfi (University of Szeged, Hungary), Valentina Mariotti (University of Bologna, Italy; Aix-Marseille Université, France), Alessandro Riga (University of Florence, Italy), M. Giovanna Belcastro (University of Bologna, Italy; Aix-Marseille Université, France), Jesper L. Boldsen (University of Southern Denmark, Denmark), and Charlotte Avanzi (Colorado State University, USA). The authors would also like to thank the laboratory team of the MPI for the Science of Human History in Jena for extensive support with capture experiments and sequencing and the developer of EAGER 2: Alexander Peltzer and James Fellows Yates (Max Planck Institute for the Science of Human History, Jena, Germany). This work was supported by the University of Zurich’s University Research Priority Program “Evolution in Action: From Genomes to Ecosystems” (V.J.S), the Max-Planck Society (J.K., A.H.), the Senckenberg Centre for Human Evolution and Palaeoenvironment (S-HEP) at the University of Tübingen (V.J.S., J.K., A.F.). P.S.’s research work is supported by ICMR, DBT India, R2STOP Canada, and the Leprosy Research Initiative Netherlands. The manuscript has been approved by the Publication Screening Committee of ICMR-NIRTH, Jabalpur, and assigned with the number ICMR-NIRTH/PSC/44/2020.

