## Supplementary Information for "Comparison of target enrichment strategies for ancient pathogen DNA"

Anja Furtwängler[1,2], Judith Neukamm[1,3,4], Lisa Böhme[5], Ella Reiter[1], Melanie Vollstedt[5], Natasha Arora[6], Pushpendra Singh[7], Stewart T. Cole[8], Sascha Knauf[9,10], Sébastien Calvignac-Spencer[11], Ben Krause-Kyora[5,12], Johannes Krause[1,2,12], Verena J. Schuenemann[1,2,3#], Alexander Herbig[1,12#]

[1]Institute for Archaeological Sciences, Archaeo- and Palaeogenetics, University of Tübingen, Germany

[2]Senckenberg Centre for Human Evolution and Palaeoenvironment, University of Tübingen, Germany

[3]Institute of Evolutionary Medicine, University of Zurich, Switzerland.

[4]Institute for Bioinformatics and Medical Informatics, University of Tübingen, Germany

[5]Institute of Clinical Molecular Biology, Kiel University, Germany

[6]Zurich Institute of Forensic Medicine, University of Zurich, Switzerland

[7]Indian Council of Medical Research-National Institute of Research in Tribal Health, Jabalpur, MP, India

[8]Institut Pasteur, Paris, France

[9]Deutsches Primatenzentrum GmbH, Leibniz-Institute for Primate Research, Goettingen, Germany

[10]Department for Animal Sciences, Georg-August-University, Goettingen, Germany

[11]Robert Koch Institut, Berlin, Germany

[12]Max Planck Institute for the Science of Human History, Jena, Germany

#These authors jointly supervised this study.

#### Supplementary Note 1 – Laboratory workflow

##### Sample Selection

The modern and ancient DNA extracts used in this study were previously tested positive for DNA from *Mycobacterium leprae* (**Schuenemann et al. 2013, Mendum et al. 2014, Schuenemann et al. 2018a**) or *Treponema pallidum* (**Arora et al. 2016, Knauf et al. 2018, Schuenemann et al. 2018b**). Existing libraries of the previous studies were used. For a comparison of the three methods under investigation, an additional shotgun sequencing of all samples was performed to determine the percentage of target DNA in the libraries prior to enrichment.

##### Ethics statement

For all samples used in this study only existing sequencing libraries were used, and no new material was collected. Statements about ethical approval and research permission can be found in the original publications (Supplementary Table 1). In this study only sequencing data of the two bacteria and no human data was generated.

##### Array capture

The array capture was performed according to the methods described in **Hodges et al. 2007**. The array design was identical to **Schuenemann et al. 2013** (*M. leprae*) and **Arora et al. 2016** (*T. pallidum*). Probe length on both arrays was 60 bp. Modern and ancient samples positive for *M. leprae* were pooled equimolar and captured on two arrays. For the *Treponema* samples, three pools for the capture were prepared: one for modern and ancient syphilis samples respectively and a third of positive extracts originating from different species of nonhuman primates. After the hybridization, the products were quantified by qPCR as described in **Schuenemann et al. 2013**. After determination of the sufficient number of cycles the pools were amplified and after quantification on an Agilent 2100 Bioanalyzer the pools were diluted for sequencing on a

HiSeq4000 using a 75bp single-end kit for the first replicates and 75bp paired-end for the following two replicates.

**In-solution capture with RNA baits (MYBaits)**

The in-solution capture was performed using biotinylated RNA baits from the MYBaits from MYcroarray® according to manufactures instructions. The first replicate was performed in the post-amplification laboratory of the Ancient DNA Laboratory at the Kiel University after the manual of version 1.3.8. The following two replicates were performed in the post-amplification laboratory of the AG Palaeogenetics at the Institute for Archaeological Science in Tübingen after the manual of version 3.02. Samples were pool identical to the array capture and the capture was followed by a similar procedure to prepare for sequencing.

For the baits for *M. leprae* the reference Br4923 (NC_011896) was used as a base for the bait design with a 2x tiling as well and 76,490 baits in the final set. Bait length was 80 bp for *M. leprae*. With regard to *T. pallidum* the baits spanned the simian derived *T. pallidum* ssp. *pertenue* strain Fribourg-Blanc reference genome (NC_021179.1) with a 2x tiling as described in **Knauf et al. 2018**. Bait length for *T. pallidum* is 100bp and the bait set contains 19,925 unique probes.

The two rounds from the first repetition were sequenced on a HiSeq4000 with a 75bp single-end kit. The two rounds of the second and third repetition using a 75bp paired-end kit.

**In-solution capture with DNA baits (probes derived from arrays)**

The in-solution capture with array probes was performed at the Max Planck Institute for the Science of Human History in Jena.

For *M. leprae,* probes were designed based on strains TN (NC_002677.1) and Br4923 (NC_011896.1). For targeted enrichment of *Treponema pallidum* DNA probes were designed on the basis of *T. pallidum* ssp. *pallidum* strains Nichols (NC_000919.1), SS14 (NC_021508.1),

Sea 81-4 (NZ_CP003679.1), Mexico A (NC_018722.1), *T. pallidum* ssp. *endemicum* strain Bosnia A (NZ_CP007548.1), and *T. pallidum* ssp. *pertenue* strain Fribourg-Blanc (NC_021179.1). The tiling density is two and one bp for *M. leprae* and *T. pallidum*, respectively. For both target organisms the probe length is 52 bp with an additional 8bp linker sequence (CACTGCGG) as described in **Fu et al. 2013**. Duplicated probes and probes with low sequence complexity were removed. This resulted in 1,125,985 and 1,593,068 unique probe sequences for *T. pallidum* and *M. leprae*, respectively. For each target species the probe set was spread on two Agilent one-million feature SureSelect DNA Capture Arrays. The capacity of the two arrays was filled by randomly duplicating probes from the probe set. The arrays were turned into in-solution DNA capture libraries as described in **Fu et al. 2013**. All three replicates were sequenced on a HiSeq4000 using a 75bp paired-end kit.

#### Supplementary Note 2 - Bioinformatics and Statistical Analysis

The sequencing data of all samples was processed with the EAGER 2 pipeline (**Peltzer et al. 2016, Fellows Yates et al. 2020,** https://github.com/nf-core/eager). Including mapping with BWA, removal of duplicates, and the generation of damage plots. The enrichment factor for all enriched libraries was calculated by dividing the percentage of endogenous DNA after enrichment by the percentage of endogenous DNA in the shotgun sequencing.

All statistical analysis was performed in R version 426 3.4.3 (R Core Team 2017). The significance between the different features mean coverage, standard deviation of the mean coverage, percentage of the genome covered five-old, fragment length and ancient DNA (aDNA) specific damage in the data of the three tested protocols was assessed in each sample individually with a mixed model as implemented in the lme4-package for R (**Bates et al. 2015**). This model contained the replicate as a random slope factor allowing between-replicate variation in the main effect. Subsequently, p-values were corrected for multiple testing with the Bonferroni correction. Pairwise Tukey HSD post hoc tests were calculated from this model using the *lsmeans* package for R (**Lenth 2016**).

Also, significant differences between the individual replicates (grouped by age and pathogen) were assessed using a linear model as implemented in the stats-package for R (R Core Team (2019)).

The percentage of unique reads was calculated by dividing the number of unique reads by the number of total reads mapped.

**Variance within each method**

After calculating the absolute value of the pairwise differences between the replicates for each method we used one-way ANOVA to determine the significance of these differences followed by Tukey HSD post hoc tests.

#### Specificity of the three tested methods

We also used one-way ANOVA to determine the significance of differences in the ratio of specific reads for each sample individually followed by Tukey HSD post hoc tests. Subsequently, p-values were corrected for multiple testing with the Bonferroni correction. General mycobacterial or treponemal reads, as well as specific reads, were determined using the MALT algorithm (**Vågene et al. 2018**). For the ratio, the number of specific reads was divided by either the number of total mycobacterial or treponemal reads.

#### Data upload

For the samples derived from human patients the reads mapping to the human genome were removed from the fastq files prior data upload with the --*strip_input_fastq* flag of EAGER 2 (**Peltzer et al. 2016, Fellows Yates et al. 2020,** https://github.com/nf-core/eager) while mapping to the hg19 reference genome.

### Supplementary Note 3 – Results of the Bioinformatics and Statistical Analysis

#### Capture efficiency

Detailed results of the tests for significant differences in the mean coverage, standard deviation of the mean coverage, percentage of the genome covered five-fold, enrichment factor, as well as aDNA typical damage and fragment length for each individual sample can be found in Supplementary Table 2 and Supplementary Figures 1 & 2.

Mean coverage and the percentage of the genome covered at least five-fold are highly dependent on the enrichment factor and therefore the results for the two features mirror that of the enrichment factor (see Main Manuscript and Supplementary Table 2).

For the features enrichment factor and mean coverage in the ancient data of both bacteria, the RNA bait capture with two rounds of hybridization shows the best results. However, adjusted p-values do not reach significance. The fragment length is in the data of ancient samples of both bacteria the shortest in the in-solution capture with DNA probes. The differences in fragment length are significant in both cases and increase with the bait length used with the longest fragments in the RNA bait capture with either one or with two rounds of hybridization.

The evenest coverage as represented by lowest values of the standard deviation of the mean coverage is seen in the DNA bait capture in the ancient *T. pallidum* samples and in the data of the RNA bait capture with two rounds of hybridization in the ancient *M. leprae* samples. In both cases, differences are not significant.

The largest portions of the genome covered at least five-fold result from RNA bait capture with two rounds of hybridization in ancient *M. leprae* sample (three to twenty times higher) and from DNA bait capture in the ancient *T. pallidum* samples (in average a hundred times higher, results of statistical significance in Supplementary Table 2). However, also in these cases the adjusted p-values do not reach significance.

An important characteristic of ancient sequencing libraries is the occurrence of the substitution of C by T at the fragment ends (**Briggs et al. 2007**). This is due to the post-mortem decay of the DNA and can be used to authenticate ancient DNA. In the data of both bacteria the array capture results in the highest portion (up to two times higher) of damaged fragment. However, differences are not significant.

Also, in the modern samples enrichment factor and mean coverage are the highest (between three and six hundred times higher) in the data of the RNA bait capture with two rounds of hybridization for both bacteria. All adjusted p-values for the enrichment factor and most of the adjusted p-values for the mean coverage are significant.

The evenest coverage in the modern data is found in the array capture data for *T. pallidum* and in the DNA bait capture data for the *M. leprae*. For *T. pallidum* these differences are significant. The percentage of the genome covered five-fold is highest for modern *M. leprae* in the RNA bait capture with two rounds of hybridization and for *T. pallidum* in the DNA bait capture. However, only for *M. leprae* differences are significant.

The longest fragment in the modern data are as well found in the methods with the longest baits. For *M. leprae* the RNA bait capture, as for the ancient sample with either one or two rounds of hybridization, results in the longest fragments. For *T. pallidum* the DNA bait capture results in the longest fragments, there the differences also reach significance.

All comparisons between the different methods were also performed between the individual replicates. Detailed results for this comparison can be found in Supplementary Table 3. The general pattern found in sample wise comparison was confirmed with higher statistical significance in the tests performed per replicate.

The number of unique reads in the data of the first and second round of hybridization with the RNA baits does not significantly increase with the second round (Supplementary Figure 3). Showing that the increase in the percentage of endogenous DNA increases while library complexity decreases.

**Variance within each method**

The choice of method significantly affects the differences between the replicates for all tested features in modern and ancient *M. leprae* genomes (Supplementary Table 6). The in-solution capture using DNA probes shows the smallest differences between the replicates besides the enrichment factor. Here the array capture produces the most similar results between the replicates (Supplementary Table 6).

For the data from the ancient and modern syphilis, the variance between the replicates is significantly affected as well. The array capture shows hereby the smallest differences between the single replicates.

**Specificity of the different methods**

The significance of differences in the ratio of specific reads for either *M. leprae* or *T. pallidum* compared to mycobacterial and treponemal reads, respectively, in each of the ancient and modern samples of both tested bacteria, was assessed (Supplementary Table 4). There is no statistical significance between the methods besides in the data of the ancient *M. leprae* samples. Here the RNA bait capture with two rounds shows significantly the highest ratios of specific reads of *M. leprae* to mycobacterial reads in total. For the ancient *Treponema pallidum* samples, the RNA bait capture with two rounds shows the highest proportions of specific reads.

For the samples of modern *M. leprae*, there is no statistical significance but here the in-solution capture with DNA probes shows the highest proportions of *M. leprae* specific reads. In the modern samples, only one round of capture with RNA baits yields the highest specificity.

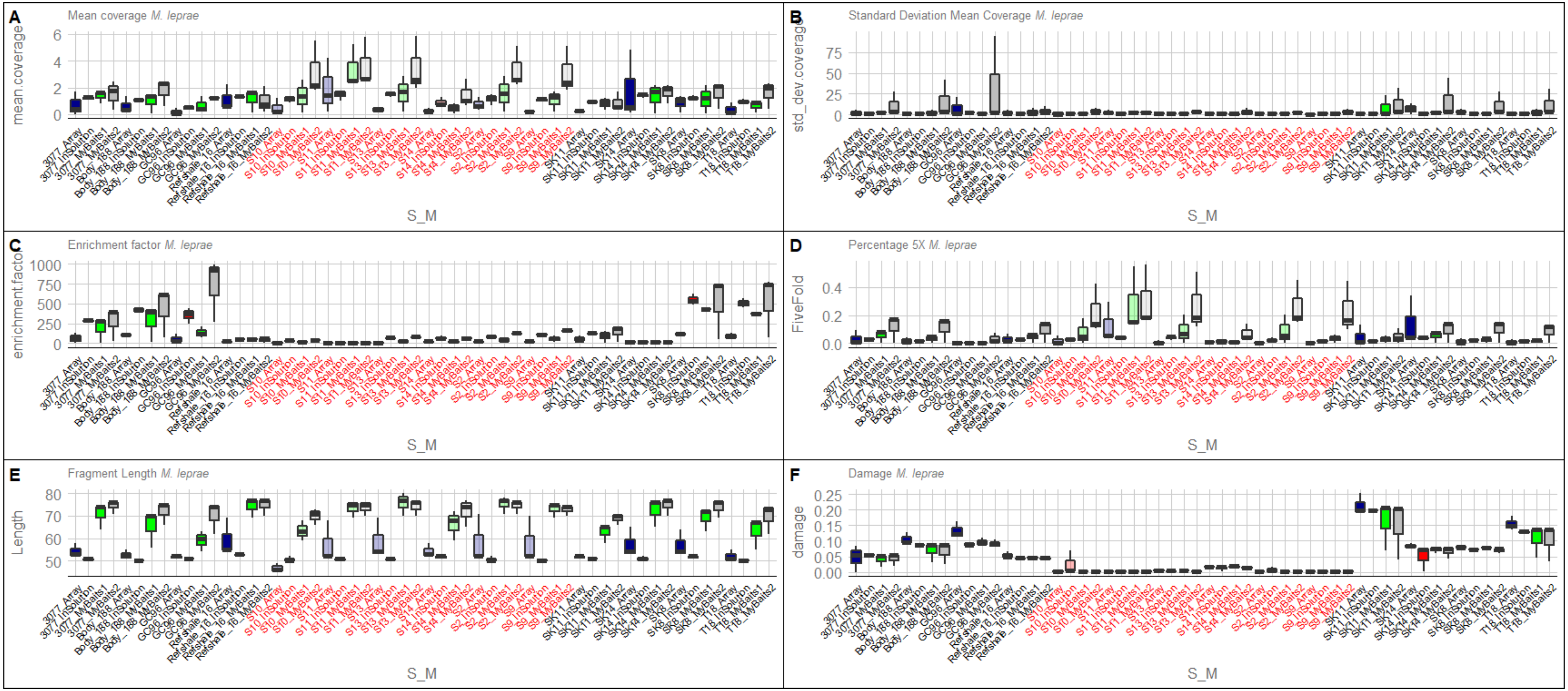


**Supplementary Figure 1: Comparison of (A) mean coverage, (B) standard deviation of the mean coverage, (C) enrichment factor, (D) and the percentage of the genome covered 5 fold, (E) distribution of the fragment length and (F) frequency of the aDNA damage for the ancient and modern strains of *M. leprae*.** Three independent replicates were performed for each method. Labels of the ancient samples are in black and for the modern samples in red. Boxplots of the array are blue, of the DNA bait capture red and the RNA baits capture is green and grey for the first and second round, respectively.

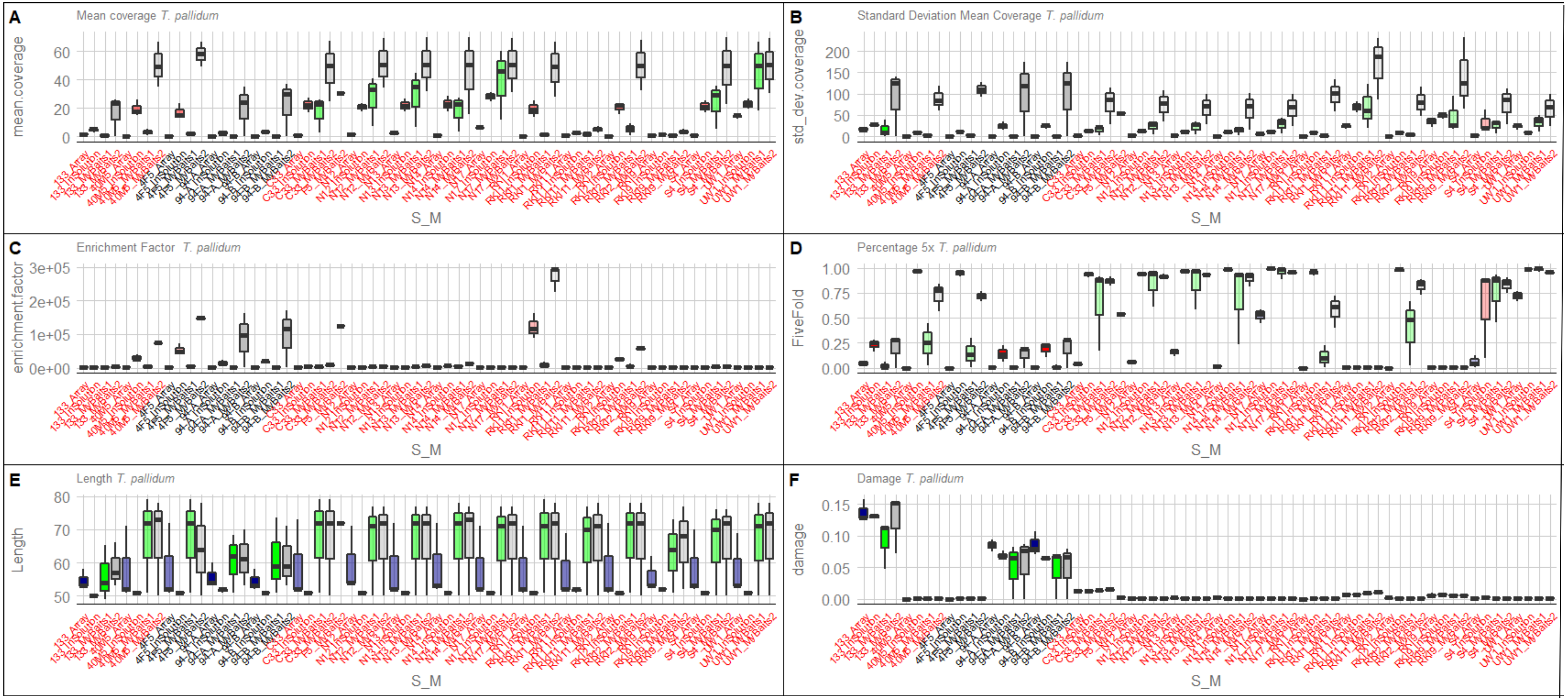


**Supplementary Figure 2: Comparison of (A) mean coverage, (B) standard deviation of the mean coverage, (C) enrichment factor, (D) and the percentage of the genome covered 5 fold, (E) distribution of the fragment length and (F) frequency of the aDNA damage for the ancient and modern strains of *T. pallidum*.** Three independent replicates were performed for each method. Labels of the ancient samples are in black and for the modern samples in red. Boxplots of the array are blue, of the DNA bait capture red and the RNA baits capture is green and grey for the first and second round, respectively.

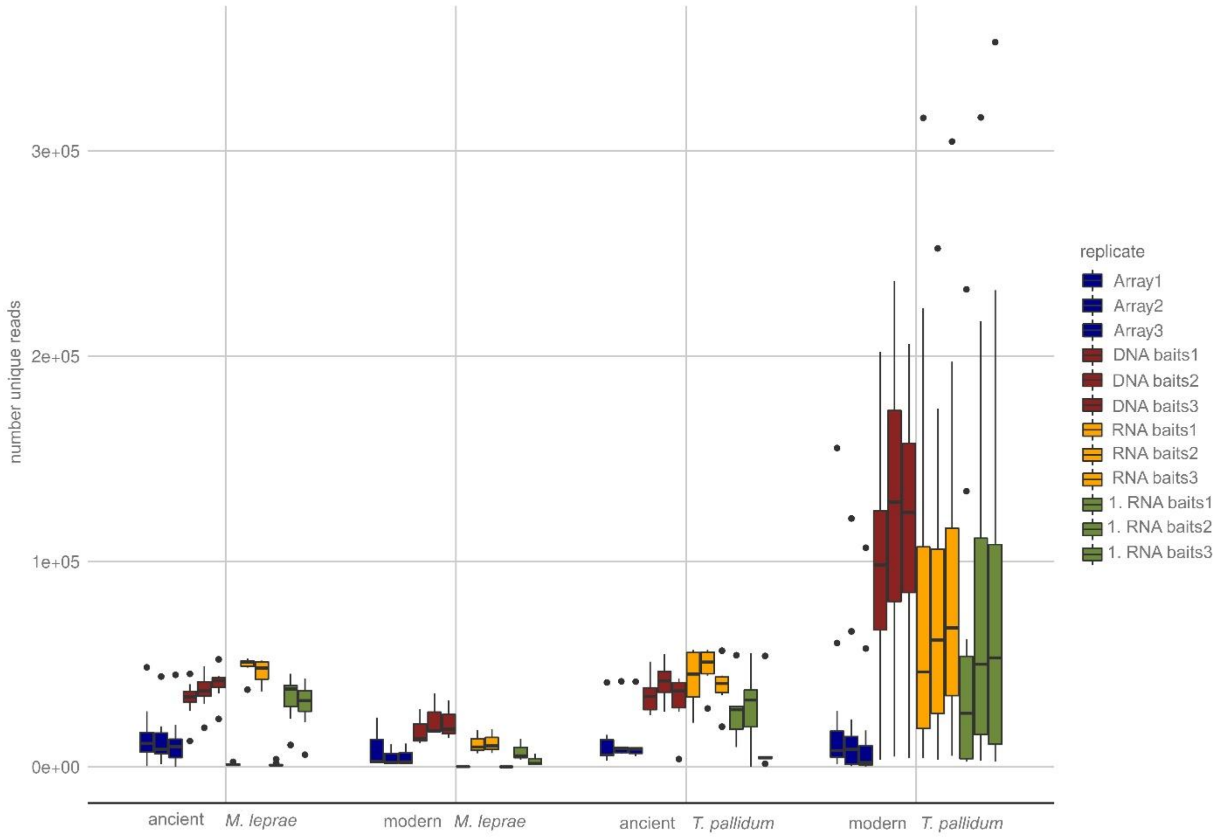


**Supplementary Figure 3: Number of unique reads for the three replicate batches of the three tested methods.** The number of unique reads in the second round of hybridization with the RNA baits does not strongly increase compared to the first round.

**Supplementary Table 1: List of all samples used in this study group according to organism and age together with the original publications.**

| Group | Sample | Age | Organism | Host species | publication |
|---|---|---|---|---|---|
| modern *T. pallidum* retrieved from nonhuman primates | 40M5160407 | modern | *T. pallidum* | nonhuman primate | Knauf et al. 2018 |
| | 4F5230307 | modern | *T. pallidum* | nonhuman primate | Knauf et al. 2018 |
| | RKI1 | modern | *T. pallidum* | nonhuman primate | Knauf et al. 2018 |
| | RKI11 | modern | *T. pallidum* | nonhuman primate | Knauf et al. 2018 |
| | RKI2 | modern | *T. pallidum* | nonhuman primate | Knauf et al. 2018 |
| | RKI9 | modern | *T. pallidum* | nonhuman primate | Knauf et al. 2018 |
| modern *T. pallidum* | N12 | modern | *T. pallidum* | human | Arora et al. 2016 |
| | N13 | modern | *T. pallidum* | human | Arora et al. 2016 |
| | N14 | modern | *T. pallidum* | human | Arora et al. 2016 |
| | N17 | modern | *T. pallidum* | human | Arora et al. 2016 |
| | C33 | modern | *T. pallidum* | human | Arora et al. 2016 |
| | S4 | modern | *T. pallidum* | human | Arora et al. 2016 |
| | UW1 | modern | *T. pallidum* | human | Arora et al. 2016 |
| Ancient *M. leprae* | 3077 | ancient | *M. leprae* | human | Schuenemann et al 2013 |
| | GC96 | ancient | *M. leprae* | human | Schuenemann et al 2018a |
| | Body_188 | ancient | *M. leprae* | human | Schuenemann et al 2018a |
| | SK11 | ancient | *M. leprae* | human | Schuenemann et al 2018a |
| | T18 | ancient | *M. leprae* | human | Schuenemann et al 2018a |
| | Refshale_16 | ancient | *M. leprae* | human | Schuenemann et al 2013 |
| | SK14 | ancient | *M. leprae* | human | Mendum et al. 2014 |
| | SK8 | ancient | *M. leprae* | human | Mendum et al. 2014 |
| modern *M. leprae* | S10 | modern | *M. leprae* | human | Schuenemann et al 2013 |
| | S11 | modern | *M. leprae* | human | Schuenemann et al 2013 |
| | S13 | modern | *M. leprae* | human | Schuenemann et al 2013 |
| | S14 | modern | *M. leprae* | human | Schuenemann et al 2013 |
| | S2 | modern | *M. leprae* | human | Schuenemann et al 2013 |
| | S9 | modern | *M. leprae* | human | Schuenemann et al 2013 |
| ancient *T. pallidum* | 94-A | ancient | *T. pallidum* | human | Schuenemann et al 2018b |
| | 133 | ancient | *T. pallidum* | human | Schuenemann et al 2018b |
| | 94-B | ancient | *T. pallidum* | human | Schuenemann et al 2018b |

#### Supplementary Table 4: Comparison of the specific reads of the three tested protocols.

| Sample | age | Organism | Chisq | Pr(>Chisq) | Array | RNA baits1 | RNA baits2 | DNA baits | Array - RNA baits1 | Array - RNA baits2 | Array - DNA baits | RNA baits1 - RNA baits2 | RNA baits1 - DNA baits | RNA baits2 - DNA baits | p.adjust |
|---|---|---|---|---|---|---|---|---|---|---|---|---|---|---|---|
| 40M5160407 | modern | T. pallidum | 1.26 | 0.35 | 0.28 | 0.33 | 0.33 | 0.28 | 0.53 | 0.55 | 1.00 | 1.00 | 0.54 | 0.56 | 1.00 |
| 4F5230307 | modern | T. pallidum | 0.73 | 0.57 | 0.30 | 0.33 | 0.32 | 0.28 | 0.85 | 0.97 | 0.93 | 0.99 | 0.54 | 0.76 | 1.00 |
| C33 | modern | T. pallidum | 1.02 | 0.43 | 0.33 | 0.35 | 0.36 | 0.30 | 0.92 | 0.91 | 0.82 | 1.00 | 0.49 | 0.47 | 1.00 |
| 94-A | ancient | T. pallidum | 1.48 | 0.29 | 0.33 | 0.35 | 0.38 | 0.31 | 0.92 | 0.50 | 0.95 | 0.84 | 0.66 | 0.26 | 1.00 |
| 133 | ancient | T. pallidum | 7.43 | 0.01 | 0.28 | 0.32 | 0.34 | 0.29 | 0.09 | 0.01 | 0.95 | 0.61 | 0.18 | 0.03 | 0.17 |
| 94-B | ancient | T. pallidum | 2.38 | 0.15 | 0.30 | 0.35 | 0.37 | 0.29 | 0.53 | 0.24 | 1.00 | 0.91 | 0.44 | 0.19 | 1.00 |
| N12 | modern | T. pallidum | 0.96 | 0.46 | 0.32 | 0.35 | 0.35 | 0.30 | 0.86 | 0.86 | 0.91 | 1.00 | 0.51 | 0.52 | 1.00 |
| N13 | modern | T. pallidum | 0.87 | 0.50 | 0.33 | 0.35 | 0.35 | 0.30 | 0.91 | 0.92 | 0.87 | 1.00 | 0.54 | 0.54 | 1.00 |
| N14 | modern | T. pallidum | 0.92 | 0.47 | 0.33 | 0.35 | 0.35 | 0.30 | 0.88 | 0.88 | 0.90 | 1.00 | 0.52 | 0.53 | 1.00 |
| N17 | modern | T. pallidum | 0.99 | 0.45 | 0.33 | 0.35 | 0.35 | 0.30 | 0.85 | 0.85 | 0.91 | 1.00 | 0.51 | 0.50 | 1.00 |
| RKI1 | modern | T. pallidum | 0.97 | 0.45 | 0.29 | 0.33 | 0.33 | 0.29 | 0.69 | 0.73 | 1.00 | 1.00 | 0.56 | 0.60 | 1.00 |
| RKI11 | modern | T. pallidum | 2.69 | 0.12 | 0.00 | 0.00 | 0.00 | 0.00 | 0.13 | 1.00 | 0.66 | 0.17 | 0.56 | 0.76 | 1.00 |
| RKI2 | modern | T. pallidum | 1.00 | 0.44 | 0.32 | 0.33 | 0.33 | 0.29 | 0.94 | 0.95 | 0.78 | 1.00 | 0.47 | 0.49 | 1.00 |
| RKI9 | modern | T. pallidum | 3.50 | 0.07 | 0.00 | 0.01 | 0.00 | 0.00 | 0.10 | 1.00 | 1.00 | 0.10 | 0.14 | 1.00 | 1.00 |
| S4 | modern | T. pallidum | 0.70 | 0.58 | 0.32 | 0.35 | 0.35 | 0.31 | 0.85 | 0.87 | 0.98 | 1.00 | 0.65 | 0.66 | 1.00 |
| UW1 | modern | T. pallidum | 1.31 | 0.34 | 0.32 | 0.36 | 0.36 | 0.30 | 0.63 | 0.65 | 0.99 | 1.00 | 0.45 | 0.47 | 1.00 |
| 3077 | ancient | M. leprae | 0.79 | 0.53 | 0.99 | 0.93 | 0.76 | 0.99 | 0.98 | 0.57 | 1.00 | 0.78 | 0.98 | 0.57 | 1.00 |
| GC96 | ancient | M. leprae | 0.37 | 0.77 | 0.80 | 0.69 | 0.68 | 0.94 | 0.98 | 0.97 | 0.95 | 1.00 | 0.81 | 0.79 | 1.00 |
| Body 188 | ancient | M. leprae | 0.54 | 0.67 | 0.95 | 0.71 | 0.68 | 0.96 | 0.84 | 0.80 | 1.00 | 1.00 | 0.82 | 0.77 | 1.00 |
| SK11 | ancient | M. leprae | 0.61 | 0.63 | 0.98 | 0.85 | 0.74 | 0.97 | 0.92 | 0.67 | 1.00 | 0.95 | 0.92 | 0.68 | 1.00 |
| T18 | ancient | M. leprae | 0.60 | 0.63 | 0.96 | 0.86 | 0.73 | 0.98 | 0.97 | 0.71 | 1.00 | 0.92 | 0.94 | 0.64 | 1.00 |
| Refshale_16 | ancient | M. leprae | 0.86 | 0.50 | 1.00 | 0.99 | 0.92 | 0.99 | 1.00 | 0.55 | 1.00 | 0.63 | 1.00 | 0.57 | 1.00 |
| SK14 | ancient | M. leprae | 0.75 | 0.55 | 0.99 | 0.97 | 0.91 | 0.99 | 0.99 | 0.59 | 1.00 | 0.78 | 0.99 | 0.59 | 1.00 |
| SK8 | ancient | M. leprae | 0.64 | 0.61 | 0.98 | 0.85 | 0.74 | 0.99 | 0.93 | 0.68 | 1.00 | 0.94 | 0.91 | 0.64 | 1.00 |
| S10 | modern | M. leprae | 6.40 | 0.02 | 0.99 | 1.00 | 1.00 | 0.99 | 0.81 | 0.17 | 0.29 | 0.51 | 0.08 | 0.01 | 0.23 |
| S11 | modern | M. leprae | 22.37 | 0.00 | 1.00 | 1.00 | 1.00 | 0.99 | 0.21 | 0.08 | 0.01 | 0.89 | 0.00 | 0.00 | 0.00 |
| S13 | modern | M. leprae | 12.10 | 0.00 | 1.00 | 1.00 | 1.00 | 0.99 | 0.75 | 1.00 | 0.00 | 0.71 | 0.01 | 0.00 | 0.03 |
| S14 | modern | M. leprae | 11.12 | 0.00 | 1.00 | 1.00 | 1.00 | 0.99 | 0.83 | 0.30 | 0.02 | 0.10 | 0.07 | 0.00 | 0.04 |
| S2 | modern | M. leprae | 13.49 | 0.00 | 1.00 | 1.00 | 1.00 | 0.99 | 0.04 | 0.08 | 0.21 | 0.94 | 0.00 | 0.00 | 0.02 |
| S9 | modern | M. leprae | 5.62 | 0.02 | 0.99 | 1.00 | 1.00 | 0.99 | 0.15 | 0.03 | 0.93 | 0.64 | 0.32 | 0.06 | 0.32 |

#### Supplementary Table 6: Comparison of the variance within each method tested.

| Organism | Feature | Sample Group | Chisq | Pr(>Chisq) | Array | DNA baits | RNA baits1 | RNA baits2 | Array - RNA baits1 | Array - RNA baits2 | Array - DNA baits | RNA baits1 - RNA baits2 | RNA baits1 - DNA baits | RNA baits2 - DNA baits |
|---|---|---|---|---|---|---|---|---|---|---|---|---|---|---|
| *M. leprae* | Mean Coverage | ancient | 19.44 | 2.21E-04 | 0.97 | 0.19 | 0.89 | 1.04 | 0.00 | 0.99 | 0.99 | 0.01 | 0.00 | 0.91 |
| | | modern | 30.28 | 1.21E-03 | 0.76 | 0.34 | 1.42 | 2.10 | 0.59 | 0.18 | 0.00 | 0.01 | 0.00 | 0.17 |
| | standard deviation Mean Coverage | ancient | 39.76 | 1.20E-05 | 4.22 | 0.10 | 4.01 | 24.20 | 0.72 | 1.00 | 0.00 | 0.75 | 0.00 | 0.00 |
| | | modern | 28.90 | 2.35E-03 | 1.15 | 0.19 | 0.95 | 2.52 | 0.10 | 0.96 | 0.01 | 0.25 | 0.00 | 0.00 |
| | Enrichment Factor | ancient | 32.09 | 5.00E-04 | 36.60 | 48.60 | 152.00 | 277.00 | 0.99 | 0.09 | 0.00 | 0.16 | 0.00 | 0.06 |
| | | modern | 30.65 | 1.01E-06 | 4.44 | 14.80 | 28.90 | 12.70 | 0.17 | 0.00 | 0.36 | 0.03 | 0.97 | 0.01 |
| | 5Xpercentage | ancient | 27.64 | 4.33E-06 | 0.57 | 0.01 | 0.03 | 0.09 | 0.01 | 0.40 | 0.17 | 0.43 | 0.00 | 0.00 |
| | | modern | 35.26 | 1.07E-04 | 0.04 | 0.02 | 0.12 | 0.21 | 0.93 | 0.10 | 0.00 | 0.02 | 0.00 | 0.06 |
| | damage | ancient | 29.31 | 1.92E-03 | 4.50 | 0.42 | 6.75 | 5.50 | 0.00 | 0.23 | 0.83 | 0.00 | 0.00 | 0.71 |
| | Length | ancient | 24.89 | 1.63E-02 | 8.78 | 0.78 | 5.78 | 4.56 | 0.00 | 0.21 | 0.04 | 0.01 | 0.07 | 0.85 |
| | | modern | 10.30 | 1.62E-02 | 0.03 | 0.01 | 0.03 | 0.04 | 0.35 | 0.95 | 0.65 | 0.13 | 0.03 | 0.92 |
| *T. pallidum* | Mean Coverage | ancient | 44.24 | 1.34E-09 | 0.20 | 1.44 | 0.68 | 21.60 | 0.97 | 1.00 | 0.00 | 0.99 | 0.00 | 0.00 |
| | | modern | 101.62 | 2.20E-16 | 0.49 | 5.02 | 13.80 | 22.80 | 0.18 | 0.00 | 0.00 | 0.00 | 0.00 | 0.00 |
| | standard deviation Mean Coverage | ancient | 45.97 | 5.76E-10 | 1.79 | 7.80 | 10.60 | 108.00 | 0.97 | 0.92 | 0.00 | 1.00 | 0.00 | 0.00 |
| | | modern | 140.84 | 2.20E-16 | 2.47 | 6.22 | 20.60 | 58.40 | 0.85 | 0.00 | 0.00 | 0.01 | 0.00 | 0 |
| | Enrichment Factor | ancient | 27.10 | 5.62E-06 | 158.00 | 5.346.00 | 1.286.00 | 74.836.00 | 0.99 | 1.00 | 0.00 | 0.99 | 0.00 | 0.00 |
| | | modern | 14.08 | 2.80E-03 | 182.00 | 7.047.00 | 2.979.00 | 5.823.00 | 0.03 | 0.68 | 0.11 | 0.37 | 0.96 | 0.67 |
| | 5Xpercentage | ancient | 23.05 | 3.95E-05 | 0.01 | 0.09 | 0.02 | 0.17 | 0.09 | 0.98 | 0.00 | 0.18 | 0.12 | 0.00 |
| | | modern | 39.93 | 1.11E-08 | 0.03 | 0.06 | 0.23 | 0.08 | 0.81 | 0.00 | 0.48 | 0.00 | 0.95 | 0.00 |
| | damage | ancient | 36.99 | 4.62E-08 | 4.00 | 0.22 | 12.50 | 10.70 | 0.20 | 0.00 | 0.01 | 0.00 | 0.00 | 0.76 |
| | Length | ancient | 101.76 | 2.20E-16 | 12.30 | 0.10 | 18.00 | 18.30 | 0.00 | 0.01 | 0.00 | 0.00 | 0.00 | 1.00 |
| | | modern | 21.26 | 9.31E-05 | 0.02 | 0.00 | 0.05 | 0.06 | 0.63 | 0.07 | 0.02 | 0.00 | 0.00 | 0.94 |

#### Supplementary Table 7: Comparison of the costs per reaction.

| in-solution capture with array derived probes | | | | | | |
|---|---|---|---|---|---|---|
| **Product** | **Supplier** | **Art.-Nr.** | **amount** | **Price (€)** | **Price (€)/rxn** | **Machines** |
| 2x Hi-RPM Hybridization Buffer | Agilent | 5190-0403 | 25 ml | 480 | 0,65 | Vortexer |
| bait production | | | 200ul | 432 | 16,20 | centrifuge |
| Herculase II Fusion DNA Polymerases | Agilent | 600679 | 400 rxn | 456 | 2,28 | centrifuge for plates |
| MinElute PCR purification Kit | Qiagen | 28004 | 50 rxn | 127 | 2,54 | Magnetic rack (96 well plate) |
| Acetic acid 100% | Sigma | 5438080100 | 100 ml | 61 | 0,00 | Magnetic rack (2 ml tubes) |
| Dynabeads MyOne T1 | ThermoFisher | 65601 | 2 ml | 546 | 10,92 | Thermocycler |
| SeraMag Speedbeads | ThermoFisher | 65152105050250 | 15 ml | 283,73 | 0,19 | Lightcycler |
| Oligonucleotide (primers, blocking oligos) | sigma | | 30 rxn | 533,5 | 17,78 | Pipettes |
| GeneAmp 10x PCR buffer | ThermoFisher | N8080006 | 1,5 ml | 195 | 3,90 | |
| Cot-1 DNA | ThermoFisher | 15279011 | 500 ug | 310 | 1,55 | |
| Denhardt’s Solution 50x | Sigma | D2532-5ML | 5 ml | 172 | 0,21 | |
| **Additional reagents** | | | | **sum/rxn** | **56,23** | |
| SDS 20 % | Sigma | 05030-500ML-F | 500 ml | 107 | | |
| SSC 20% | ThermoFisher | AM9763 | 1 L | 101 | | |
| 0.5 M EDTA | Sigma | 324506-100ML | 100 ml | 45 | | |
| HPLC H2O | Sigma | 270733-1L | 1 L | 24 | | |
| 5 M NaCl | Sigma | S5150-1L | 1 L | 68,3 | | |
| 1 M Tris-HCl, pH 8 | ThermoFisher | AM9855G | 100 ml | 57,25 | | |
| Tween-20 100% | Sigma | P1379-25ML | 25 ml | 13 | | |
| 1 M NaOH | Applichem | A6579,1000 | 1 L | 27,2 | | |
| 3 M Sodium acetate pH 5.2 | Sigma | S7899-100ML | 100 ml | 32,4 | | |
| PM buffer | Qiagen | 19083 | 500 ml | 63,6 | | |
| Salmon SpermDNA | ThermoFisher | 15632011 | 5 ml | 133 | | |
| EtOH | Merck | 1009832511 | 2,5 L | 86 | | |

| Array | | | | | | |
|---|---|---|---|---|---|---|
| **Product** | **Supplier** | **Art.-Nr.** | **amount** | **Price (€)** | **Price (€)/rxn** | **Machines** |
| Oligo aCGH/ChIP-on-chip Hybridization Kit | Agilent | 5188-5220 | Blocking Agent + 2x Hi-RPM hybridization buffer 25 rxn | 461 | 18,44 | Vortexer |
| blocking oligos | sigma | BO4, BO6, BO8, BO10 | | 304,64 | 10,15 | Thermoshaker |
| BD Plastipak | Fisher Scientific | 10205194 | 100 x | 61,38 | 0,61 | Thermocycler |
| Hybridization Gasket Slide Kit | Agilent | G2534-60005 | 100 | 2036 | 20,36 | Hybridization Oven |
| SureSelect DNA Capture Array 1 M | Agilent | G3358A | 1 | 613 | 613,00 | 1 bigger glass bowl |
| Microarray Wash Buffer Kit | Agilent | 5188-5222 | WB 1, WB 2 40 rxn | 208 | 5,20 | 2 smaller glass bowls |
| Herculase II Fusion DNA Polymerases | Agilent | 600679 | 400 rxn | 456 | 2,28 | Array chamber |
| MinElute PCR purification Kit | Qiagen | 28004 | 50 rxn | 127 | 2,54 | centrifuge |
| **Addtional ragents** | | | | **sum/rxn** | **672,59** | Magnetic mixer |
| HPLC H2O | Sigma | 270733-1L | 1 L | 24 | | Waterbath |
| | | | | | | tweezers |
| | | | | | | rack for slides |
| | | | | | | pipettes |
| | | | | | | Lightcycler |

| MyBait | | | | | | |
|---|---|---|---|---|---|---|
| **Product** | **Supplier** | **Art.-Nr.** | **amount** | **Price (€)** | **Price (€)/rxn** | **Machines** |
| MyBait Kit | | | 48 rxn | 5000 | 104,17 | Vortexer |
| Herculase II Fusion DNA Polymerases | Agilent | 600679 | 400 rxn | 456 | 2,28 | centrifuge |
| MinElute PCR purification Kit | Qiagen | 28004 | 50 rxn | 127 | 2,54 | centrifuge for plates |
| **Additional reagents** | | | | **sum/rxn** | **108,99** | Magnetic rack (96 well plate) |
| 1 M Tris-HCl, pH 8 | ThermoFisher | AM9855G | 100 ml | 57,25 | | Magnetic rack (2 ml tubes) |
| Tween-20 100% | Sigma | P1379-25ML | 25 ml | 13 | | Thermocycler |
| HPLC H2O | Sigma | 270733-1L | 1 L | 24 | | Lightcycler |
| | | | | | | Pipettes |
